# The first European woolly rhinoceros mitogenomes retrieved from within cave hyena coprolites suggest long-lasting phylogeographic differentiation

**DOI:** 10.1101/2023.06.23.546246

**Authors:** PA Seeber, Z Palmer, A Schmidt, A Chagas, K Kitagawa, E Marinova-Wolff, Y Tafelmaier, LS Epp

**Author notes:** Lead contact: LS Epp.

## Abstract

The woolly rhinoceros (*Coelodonta antiquitatis*) is an iconic species of the Eurasian Pleistocene megafauna, which was abundant in Eurasia in the Pleistocene until its demise beginning approximately 10,000 years ago. Despite the early recovery of several specimens from well-known European archaeological sites, including its type specimen (Blumenbach 1799), no genomes of European populations were available so far, and all available genomic data originated exclusively from the Siberian population^1^. Using coprolites of cave hyenas (*Crocuta crocuta spelea*) recovered from Middle Palaeolithic layers of two caves in Germany (Bockstein-Loch and Hohlenstein-Stadel), we isolated and enriched predator and prey DNA to assemble the first European woolly rhinoceros mitogenomes, in addition to cave hyena mitogenomes. These mitogenomes of European woolly rhinoceros are genetically distinct from the Siberian woolly rhinoceros, and analyses of the more complete mitogenome suggests a split of the populations potentially coinciding with the earliest fossil records of wooly rhinoceros in Europe.

## Main text

Reconstructing past ecosystems and identifying changes in genomes of extinct populations through ancient DNA (aDNA) may help retrace population dynamics, and, phylogeography and evolutionary changes. The genomic Pleistocene macro- and megafauna record of central Europe is, to date, limited, in part due to less suitable conditions for long-term DNA preservation in the environment, as opposed to e.g. permafrost. Predator coprolites preserved in caves are a valuable source for gathering genomic information on the individual predators and their prey. The extinct cave hyena (*Crocuta crocuta spelaea*) was a common predator in Europe during the Pleistocene until its extirpation approximately 14k years ago^2^, and its prey included large herbivores such as the woolly rhinoceros (*Coelodonta antiquitatis*), as evidenced by numerous macrofossil findings in European caves^3^. The wooly rhinoceros was a cold-adapted megaherbivore, which was abundant from western Europe to north-east Siberia during the Middle to Late Pleistocene^4^. Its fossil history suggests that, originating north of the Himalayan-Tibetan uplift around 2.5 Myr BP, *Coelodonta* spread westwards to enter Europe in the particularly cold and arid conditions of MIS12. The earliest immigration of *Coelodonta* into Eastern and Central Europe is documented by a number of finds dated to approximately 460–400 kyr BP^4,5^ from specimens that are morphologically distinct from Late Pleistocene *C. antiquitatis*. Initially assigned to a separate species, C. *tologoijensis*^*4*^, recent phylogenetic analyses of morphological characters implies inclusion in a subspecies, *C. antiquitatis praecursor*^*6*^. Irrespective of this placement, the temporal and spatial distribution of the fossils, and their morphological changes across the Middle and Late Pleistocene suggests repeated range expansions and immigration of *Coelodonta* into Central and Western Europe during successive cold periods. However, despite the wide distribution of this species throughout northern Eurasia and numerous findings of remains in western Europe, comparably little genomic information is available to investigate this. All currently published mitogenomic data of wooly rhinoceroses stem from Siberian findings^1^, whereas no mitogenome assemblies of European woolly rhinos are available to date, and most molecular genetic studies on wooly rhinoceroses from Europe were restricted to few short markers^7^. We extracted ancient DNA from two cave hyena coprolites (referred to as HST3168 and BSVK22) retrieved from Middle Palaeolithic layers of two cave sites in the Swabian Jura (Hohlenstein-Stadel and Bockstein-Loch), Germany, and we used hybridization capture to enrich mammalian mitogenomic DNA^8^ (see Supplement). From this, we assembled the mitogenomes of wooly rhinoceroses and cave hyenas.

The enriched coprolite libraries produced 91,873,020 and 61,090,867 raw reads, respectively. Reads assigned to taxa at a higher rank than genus, as well as reads assigned to a taxon with fewer than 1000 reads in total were disregarded, and the remaining reads (31,577 and 445,355, respectively) were exclusively assigned to *Crocuta crocuta* and *Coelodonta antiquitatis* (Table 1). The length distribution of these reads indicated considerable fragmentation, with 39 bp average length in HST3168 and 40 bp in BSVK22. The assembled mitogenomes covered 27% of the mitochondrial genome for both species in the HST3168 library and 59% (*C. crocuta*) and 81% (*C. antiquitatis*) in the BSVK22 library. aDNA damage patterns were in accordance with the expected patterns of DNA degradation in terms high proportions of C>T transitions at the 5’- and G>A transitions at the 3’-ends (Supplemental Figure 1). *D. sumatrensis* is the closest extant relative of *C. antiquitatis*; compared with respective modern mitogenomes, the retrieved ancient mitogenomes showed shifts in nucleotide composition with higher proportions of G and T bases, and lower proportions of A and C. aDNA damage patterns showed the expected pattern regarding the *C. crocuta* mitogenomes, whereas this result was somewhat less consistent for the *C. antiquitatis* mitogenomes. This discrepancy is likely due to the markedly higher divergence of *C. antiquitatis* from the modern reference *D. sumatrensis*, compared to the cave hyena and its conspecific modern reference. Cytosine deamination due to aDNA degradation (i.e., high C>T substitution frequencies) indicates that DNA molecules are indeed ancient.

Considering such DNA decay-mediated substitutions, phylogenetic results must be interpreted with caution; however, integrating presumed aDNA decay-mediated substitutions in phylogenetic models may suggest misleading results^9,10^. As the phylogenies produced here might be confounded by this, and the mitochondrial genome of the specimen from the Hohlenstein-Stadel cave is highly fragmented, we conducted further analyses only on the more complete mitogenome from the Bockstein site. Based on Bayesian inference conducted identically to a previous study^1^, and including those sequences, the Bockstein sequence is substantially divergent from the previously published Siberian sequences, which were grouped into a common clade. The divergence time estimate of the European sequence was between 670 ka BP and 120 ka BP, while the two recovered Siberian clades split at a substantially younger time (Fig. 1). This high divergence and the inferred timing of the split suggests that the *C. antiquitatis* mitochondrial genome from Bockstein has been separated for a very long time from Siberian populations, which in contrast do not seem to display long-lasting phylogeographic patterns.^7^ This contrasts with the hypothesis of repeated range expansions into Western Europe during cold stages of the late Pleistocene, at least for the mitochondrial lineage of our sample.

**Fig. 1.**
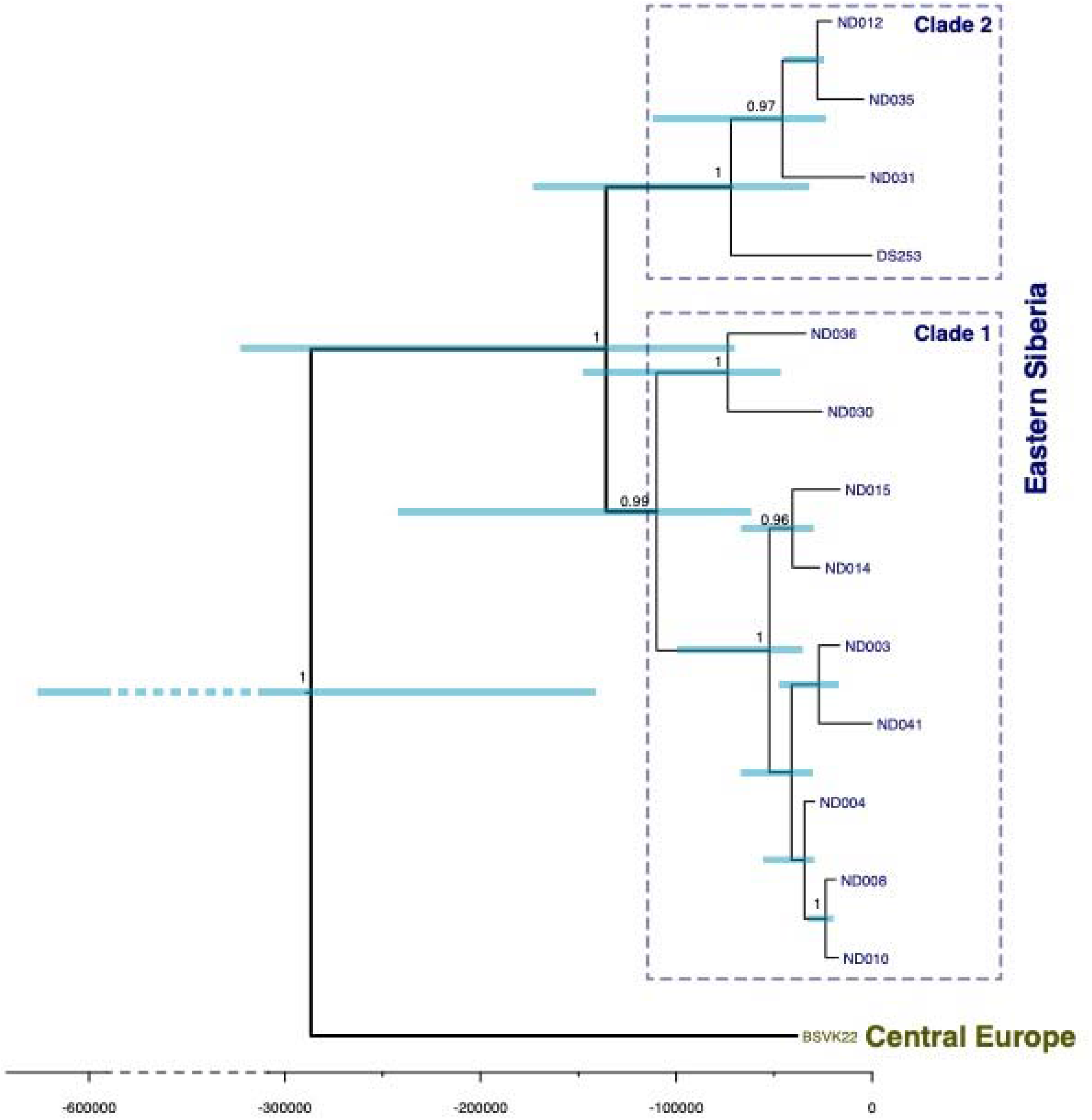
Bayesian phylogeny of the sequence from Bockstein-Loch (BSVK22) in relation to previously published sequences from Eastern Siberia^1^. Support values >0.95 are shown.

As this result is only based on a single sample from a hyena coprolite, we refrain from any further interpretation. However, the mitogenome assemblies produced here are the first mitogenomic records of European woolly rhinoceros and are thus an important resource to help resolve the phylogeography of this iconic Pleistocene megafauna species. The fact that these could be retrieved with relative ease from a coprolite of another species, i.e. no remain associated directly to woolly rhinoceros was needed, stresses the value of obtaining genomic data from a wide range of materials. As with these samples, many archaeological objects retrieved in past excavations and existing in collections, are a to date largely overlooked source of ancient DNA.

## Data availability

Raw sequence data of the hybridization capture were made available as an NCBI BioProject (PRJNA933601). Mitogenome assemblies were made available on Figshare (https://doi.org/10.6084/m9.figshare.22144169.v1).

## Acknowledgements

This research was funded through the 2017-2018 Belmont Forum and BiodivERsA joint call for research proposals, under the BiodivScen ERA-Net COFUND program, and with the funding organizations Deutsche Forschungsgemeinschaft (DFG grant EP-98/3-1 to L.S.E.), Agence Nationale de la Recherche (ANR), Research Council of Norway (NFR), the Swedish Research Council for Environment, Agricultural Sciences and Spatial Planning (Formas), Academy of Finland, National Science Foundation (NSF) and the Natural Sciences and Engineering Research Council of Canada (NSERC-CRSNG). Z.P. was supported through the CPYX - Congress-Bundestag Youth Exchange (young professionals). We thank the Museum Ulm for providing the Bockstein specimen.

