## Supplement for "The first European woolly rhinoceros mitogenomes retrieved from within cave hyena coprolites suggest long-lasting phylogeographic differentiation"

**Supplemental Experimental Procedures**

*Sites*

The two coprolites (termed HST3168 and BSVK22, respectively) derive from late Pleistocene sediments of the Palaeolithic cave sites Hohlenstein-Stadel (HST)^1^ and Bockstein-Loch (BS)^2,3^ in the Lone valley 30 km northeast of Ulm/ Germany. Both belong to the world heritage site “Caves and Ice Age Art of the Swabian Jura” and play an important role within discussions on the origin of art and music and the transition from the Middle to the Upper Palaeolithic. Both sampled objects are associated with late Middle Palaeolithic assemblages. At both sites, dental remains dominate among skeletal elements of woolly rhino, which limit the scope of taphonomic analysis.^4,5^

*DNA extraction*

Coprolite material for DNA isolation was sampled after scraping off the coprolite’s outer layer two successive times using new sterile scalpel blades each time. A dendrocorer (2 mm inner diameter) was used to retrieve material from within each coprolite. Before sampling and between coprolites, surfaces and tools were cleaned using DNA-ExitusPlus (AppliChem, Darmstadt, Germany) and ultrapure water; tools were additionally dipped in ethanol and were flamed. The coprolite samples were placed in sterile Eppendorf tubes and were transported to a dedicated aDNA laboratory at the University of Konstanz, which was physically separated from buildings containing other DNA labs. All usual precautionary measures were met to prevent contamination of aDNA; prior to these coprolite samples, the laboratory had only been used to process sedimentary ancient DNA, and no direct organismal samples had been processed in this laboratory. DNA was extracted according to (Hagan et al. 2020; method "D")^6^, with some modifications. Sample material of each coprolite (113 mg of HST3168 and 104 mg of BSVK22) was placed in 400 µL EDTA (0.5 M), and 100 µL Proteinase K (Qiagen, Hilden, Germany) was added. The suspension was placed in a PowerBead tube (Qiagen) containing 750 µL PowerBead solution and was incubated under slow rotation at room temperature for 4 h. Thereafter, bead-beating was performed twice for 30 s using a FastPrep device (MP Biomedicals, Santa Ana, CA, USA). The mixture was centrifuged at 3,400 rcf for 5 min, and the supernatant was placed in a large tube and was mixed with 15 mL Qiagen buffer PB. This mixture was transferred to the spin column of a High Pure Viral Nucleic Acid Large Volume Kit (Roche, Basel, Switzerland) for centrifugation at 1,500 rcf for 4 min, followed by rotation of each tube by 90° and centrifugation for 2 min. The silica membrane was then transferred to a sterile 1.5-mL tube and was washed twice using 700 mL buffer PE (Qiagen) under centrifugation (9,400 rcf, 1 min) and was dried (9,400 rcf, 1 min). DNA was eluted by adding 90 µL buffer EB (Qiagen) in three steps of 30 µL, each, with 5 min incubation each time. An extraction blank was processed with the samples to control for contamination. The coprolite samples yielded 72 and 81 ng DNA, respectively. After extraction (i.e., for library prep and hybridization capture), coprolite DNA was processed together with lake sedimentary DNA samples of a different study (not older than approximately 2,000 years).

*Library preparation*

Genomic libraries were produced as described previously^7^, with some modifications, using New England Biolabs reagent kits for Illumina libraries (New England Biolabs, Ipswitch, MA, USA; and Qiagen (Hilden, Germany) and were visualized through gel electrophoresis. Libraries were produced from the two DNA extracts, the extraction blank, and a library blank to control for contamination. Indexing was performed using individual combinations of P5/P7 index primers for Illumina sequencing (Illumina, San Diego, CA, USA). After indexing, libraries were pooled (four libraries per pool, together with libraries produced for a different study) and were purified using a MinElute PCR purification kit (Qiagen; elution: 2 x 20 µL). Concentrations were measured using a Qbit device (Thermo Fisher Scientific, Waltham, MA, USA).

*Hybridization capture and sequencing*

Hybridization capture was performed using RNA oligonucleotide baits (Daicel Arbor Biosciences, Ann Arbor, MI, USA); which were designed using reference sequences of 17 selected extant and extinct mammal species (Table S1; comprising 282,168 nucleotides); this set of reference sequences was compiled for a different study and did not include carnivorans^8^). The compiled mitochondrial sequences were submitted to Daicel Arbor Biosciences for custom design of 70-bp baits at three-fold tiling, collapsed at 99% identity and 85% overlap, resulting in 9,339 unique baits (747,120 nt). Libraries in pools of four were subjected to the hybridization capture reaction (the two coprolite libraries were enriched in one reaction pool together with two sediment aDNA libraries of a different study^8^). Libraries produced from extraction blanks, library blanks, and PCR non-template controls were pooled in one single reaction due to the minute DNA concentrations (all < 0.3 ng/µL, and mostly below detection range). Hybridization capture was performed according to the MYbaits protocol v. 5.00 (Daicel Arbor Biosciences) with the following modifications: 150 ng baits were used per capture reaction, and library pools were incubated for hybridization at 58 °C for 24 h. Thereafter, the enriched libraries were removed from the beads and were amplified with primers IS5/IS6 (Illumina). PCR products were purified using the MinElute PCR Purification Kit (Qiagen), and the final library concentration was measured on a Bioanalyzer (Agilent, Santa Clara, CA, USA). Enriched libraries were pooled at equal concentrations and were sequenced on an Illumina NovaSeq platform (Illumina) with an SP Flow Cell (2 x 150 bp paired end).

**Table S1.** Mitogenome templates of 17 mammal species for design of RNA baits.

| **Order** | **Species** | **NCBI accession** |
| --- | --- | --- |
| Artiodactyla | *Bison bison* | NC_012346.1 |
|  | *Bos primigenius* | NC_020746.1 |
|  | *Saiga tatarica* | NC_013996.1 |
|  | *Ovis canadensis* | NC_015889.1 |
|  | *Ovibos moschatus* | NC_020631.1 |
|  | *Cervus elaphus* | NC_007704.2 |
|  | *Rangifer tarandus* | NC_007703.1 |
|  | *Alces alces* | NC_020677.1 |
|  | *Camelus ferus* | NC_009629.2 |
| Perissodactyla | *Equus przewalskii* | NC_024030.1 |
|  | *Coelodonta antiquitatis* | NC_012681.1 |
| Lagomorpha | *Lepus arcticus* | NC_044769.1 |
|  | *Ochotona collaris* | NC_003033.1 |
| Proboscidea | *Mammuthus primigenius* | NC_007596.2 |
| Eulipotyphla | *Sorex tundrensis* | NC_025327.1 |
| Rodentia | *Castor canadensis* | NC_033912.1 |
|  | *Dicrostonyx torquatus* | NC_034646.1 |

*Data processing*

Raw sequence data were demultiplexed with bcl2fastq 2.20 (Illumina). Adapter sequences were removed and sequences were filtered using leeHom^9^. Duplicates were removed, and filtered reads were mapped against a database of 75 mammal mitogenomes (Table S2) using BWA v. 0.7.17^10^ and were blastn-aligned against the complete NCBI Genbank nucleotide database^11^; retrieved March 14 2022) with a maximum e-value of 0.01. All blast output files were processed using MEGAN Community Edition 6.19.8 using a weighted LCA algorithm^12^. aDNA degradation was examined using mapDamage 2.2.1^13^ with modern reference genomes (*Crocuta crocuta*: CM018432.1; *Dicerorhinus sumatrensis*: MF066642.1), and nucleotide composition was assessed using BBMap 38.73^14^. The mitogenomes of both species were assembled with Geneious Prime 2022.1.1^15^ using respective reference genomes downloaded from Genbank (*C. crocuta*: NC_020670.1; *C. antiquitatis*: MK909152.1).

**Table S2**. Reference mitogenomes for mapping

| **Genbank accession** | **species** | **Genbank accession** | **species** |
| --- | --- | --- | --- |
| NC_020679.1 | *Antilocapra americana* | NC_011116.1 | *Arctodus simus* |
| NC_012346.1 | *Bison bison* | NC_027963.1 | *Sorex araneus* |
| NC_020746.1 | *Saiga tatarica* | NC_025327.1 | *Sorex tundrensis* |
| NC_013996.1 | *Bos primigenius* | KJ397607.1 | *Lepus arcticus* |
| NC_015889.1 | *Ovis canadensis* | NC_001640.1 | *Equus caballus* |
| NC_020630.1 | *Oreamnos americanus* | NC_024030.1 | *Equus przewalskii* |
| NC_020631.1 | *Ovibos moschatus* | NC_018783.1 | *Equus ovodovi* |
| NC_027233.1 | *Bison priscus* | HM118851.1 | *Equus hemionus* |
| NC_009629.2 | *Camelus ferus* | MK982180.1 | *Equus asinus* |
| KR822422.1 | *Camelops cf. hesternus* | NC_012681.1 | *Coelodonta antiquitatis* |
| NC_013836.1 | *Cervus elaphus xanthopygus* | NC_009574.1 | *Mammut americanum* |
| NC_007704.2 | *Cervus elaphus* | NC_007596.2 | *Mammuthus primigenius* |
| NC_013840.1 | *Cervus elaphus yarkandensis* | FR691686.1 | *Castor fiber* |
| KP405229.1 | *Alces alces cameloides* | NC_033912.1 | *Castor canadensis* |
| NC_020677.1 | *Alces alces* | NC_034313.1 | *Dicrostonyx groenlandicus* |
| NC_020729.1 | *Odocoileus hemionus* | NC_034646.1 | *Dicrostonyx torquatus* |
| NC_015247.1 | *Odocoileus virginianus* | JN181159.1 | *Peromyscus leucopus* |
| NC_007703.1 | *Rangifer tarandus* | NC_006853.1 | *Bos taurus* |
| KY987554.1 | *Platygonus compressus* | KM093871.1 | *Capra hircus* |
| NC_002008.4 | *Canis lupus familiaris* | NC_015241.1 | *Microtus fortis fortis* |
| NC_009686.1 | *Canis lupus lupus* | KP200876.1 | *Vulpes lagopus* |
| NC_013445.1 | *Cuon alpinus* | HM236180.1 | *Ovis aries* |
| NC_026529.1 | *Vulpes lagopus* | KT448275.1 | *Canis latrans* |
| NC_028302.1 | *Panthera leo* | JN632610.1 | *Capreolus capreolus* |
| NC_022842.1 | *Panthera onca* | KJ681493.1 | *Capreolus pygargus* |
| NC_010642.1 | *Panthera tigris* | JN632629.1 | *Dama dama* |
| NC_014456.1 | *Lynx rufus* | DQ316069.1 | *Loxodonta africana* |
| NC_020642.1 | *Martes americana* | KM982549.1 | *Lynx lynx* |
| NC_020641.1 | *Neovison vison* | KP202265.1 | *Panthera pardus* |
| NC_024942.1 | *Mustela nigripes* | NC_026460.1 | *Rhinolophus macrotis* |
| NC_020664.1 | *Martes pennanti* | Y07726.1 | *Ceratotherium simum* |
| NC_020639.1 | *Mustela nivalis* | NC_005089.1 | *Mus musculus* |
| NC_009685.1 | *Gulo gulo* | AM711900.1 | *Meles meles* |
| NC_011112.1 | *Ursus spelaeus* | KM091450.1 | *Mustela erminea* |
| NC_003426.1 | *Ursus americanus* | NC_005358.1 | *Ochotona princeps* |
| NC_003427.1 | *Ursus arctos* | NC_012095.1 | *Sus scrofa domesticus* |
| NC_003428.1 | *Ursus maritimus* | DQ480489.1 | *Canis lupus familiaris* |
|  |  | NC_020670.1 | *Crocuta crocuta* |

*Mitogenome data processing*

The assembled mitogenome of C. antiquitatis stemming from Bockstein was aligned to mitogenome sequences published previously.^16^ A nearly complete set of these mitogenomes, as assembled consensus sequences, was downloaded from GenBank. These sequences and the Bockstein sequence were aligned with MUSCLE in MEGA 11^17^ and exported as a FASTA file to perform demographic reconstruction using BEAST v1.10.4^18^ equivalently to the analyses performed by Lord et al.10. The Bockstein’s sequence’s estimated age was MIS3 (45–60 kyr), and further settings, added in BEAUti v1.10.4^18^, were identical to those used by Lord et al.^16^ Tip dates of the sequences were included as years before present. The evolutionary model was HKY+I. A strict molecular clock was applied, and the clock was set to a normal distribution with the initial value as 6.1^-9^ substitutions/site/year, the mean value at 6.1^-9^ and a standard deviation of 0.01. A coalescent:constant size tree model was applied. The model was run for ten million generations with sampling every 1,000 generations. A marginal likelihood estimation of path sampling/stepping stone sampling was performed. All output log files were visualized in Tracer v1.7.1^19^ to ensure convergence had occurred. Tree Annotator v1.10.4 was used to remove 10% burn-in from the tree files, and the phylogenies were then visualized in Figtree v1.4.4.

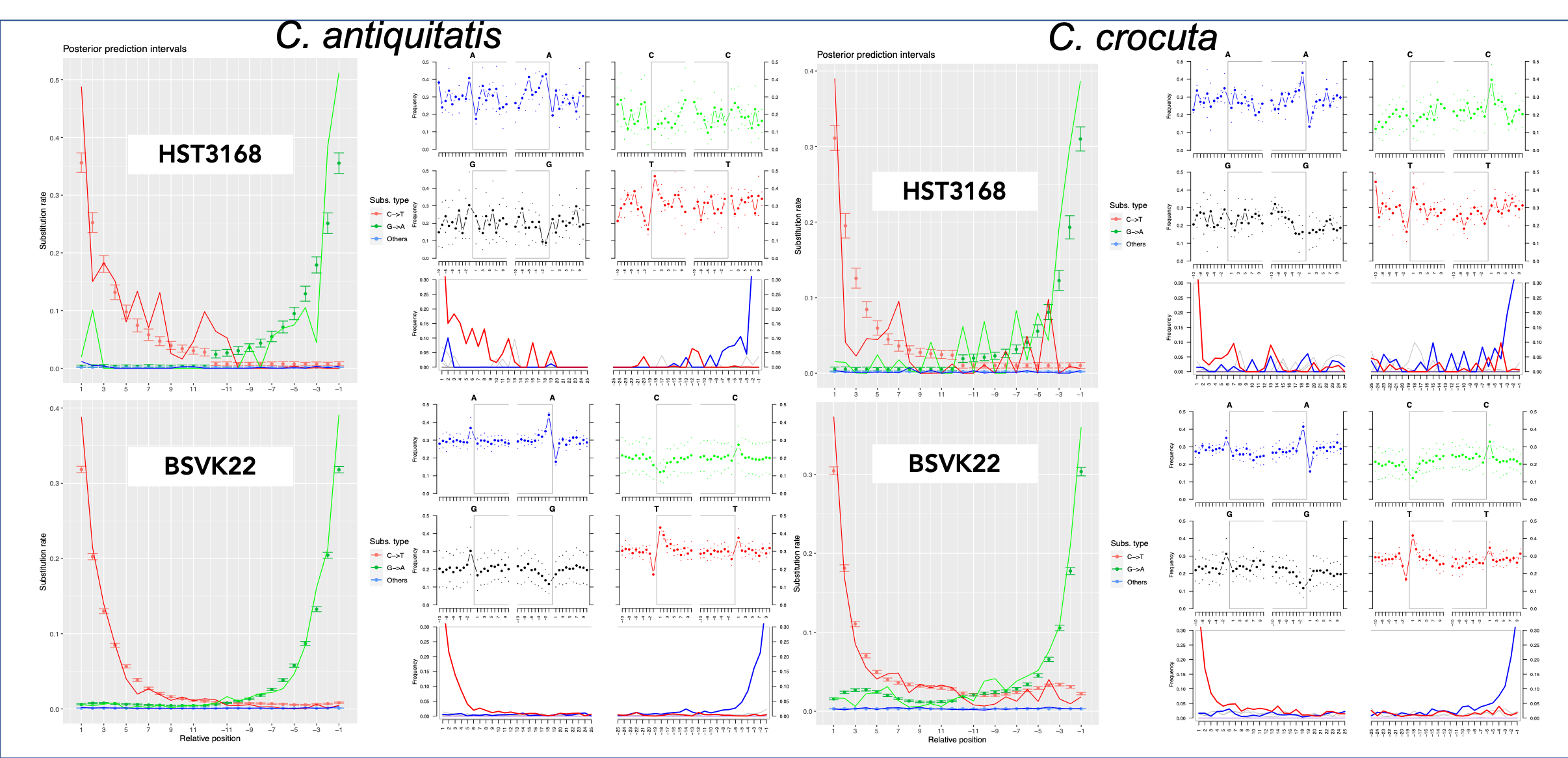

Supplemental Figure 1. aDNA damage patterns in the assembled genomes of *C. antiquitatis* and *C. crocuta* retrieved from cave hyena coprolites from Hohlenstein-Stadel (HST3168) and Bockstein-Loch (BSVK22).
